# Functional Modification of Cyanobacterial Phycobiliprotein and Phycobilisomes through Bilin Metabolism Control

**DOI:** 10.1101/2024.02.04.578791

**Authors:** Mizuho Sato, Takeshi Kawaguchi, Kaisei Maeda, Mai Watanabe, Masahiko Ikeuchi, Rei Narikawa, Satoru Watanabe

**Author notes:** **Corresponding Author** Satoru Watanabe - Department of Bioscience, Tokyo University of Agriculture, Tokyo 156-8502, Japan. **First authorship:** These authors share first authorship (Mizuho Sato, Takeshi Kawaguchi).

## Abstract

The light-harvesting antenna complexes in cyanobacteria are known as phycobilisomes (PBSs), and they can adapt to a diverse range of light environments owing to the deployment of different phycobiliproteins within the PBS structures. Freshwater cyanobacteria such as *Synechococcus elongatus* PCC 7942 thrive under red light due to phycocyanin (PC), along with phycocyanobilin (PCB), in PBS. Conversely, cyanobacteria in shorter-wavelength light environments, such as green light, employ phycoerythrin paired with phycoerythrobilin (PEB) alongside PC in PBS. Synthetic biology studies have shown that PEB production can be enhanced by additional expression of the heterologous PEB synthases, namely 15,16-dihydrobiliverdin:ferredoxin oxidoreductase and PEB:ferredoxin oxidoreductase (PebAB), leading to cellular browning due to PEB accumulation. This approach is genetically unstable, and the properties of the resulting PEB-bound PBS complexes remain uncharacterized. Herein, we engineered a novel strain of *Synechococcus* 7942 PEB1, with finely tuned control of PEB metabolism. PEB1 exhibited a remarkable and reversible color shift from green to brown and pink, based on the PebAB induction levels. High induction led to a complete PCB-to-PEB substitution, causing the disassembly of the PBS rod complex. In contrast, low PebAB levels resulted in the formation of a stable chimeric PBS complex with partial PCB-to-PEB substitution. This adaptation enabled efficient light harvesting in the green spectrum and energy transfer to the photosynthetic reaction center. These findings, which improve our understanding of PBS and highlight the structural importance of the bilin composition, lay the foundation for future PBS adaptation studies in bioengineering, synthetic biology, and renewable energy.

## INTRODUCTION

Photosynthesis, the process by which light energy is converted into chemical energy, is fundamental to sustaining life on the Earth. During evolution, photosynthetic organisms have developed various optimized light-harvesting complexes to efficiently harvest light and transfer energy to photosynthetic reaction centers in their respective environments. Phycobilisomes (PBSs), found in cyanobacteria, eukaryotic red algae, and glaucophytes, are peripheral light-harvesting complexes in the thylakoid membrane. PBS enhances the efficiency of light harvesting by channeling light energy to reaction centers for subsequent conversion into chemical energy ^1, 2^.

Structurally, PBSs consist of rod and core subcomplexes. The rods are radially attached to the core proteins. Several linear tetrapyrroles, bilins, function as chromophores and are bound to apoproteins within the rod and core subcomplexes. The rods comprise disk-like trimers, namely (αβ)_3_, which may further assemble as a hexamer ([αβ]_3_)_2_ composed of several types of phycobiliproteins, such as phycocyanins (PCs), phycoerythrins (PEs) and phycoerythrocyanins (PECs) ^3-5^. The phycobiliproteins contain covalently bound bilins, and in the case of PCs, nine phycocyanobilins (PCBs) form disk-like trimers, namely (αβ)_3_. The rods vary depending on the organism and can be made of PC only or a combination of PC and other phycobiliproteins such as PEs and PECs. The PBS core consists of a cylindrical structure composed of allophycocyanin (APC) with six PCBs per monomer. All units (disks within the rod and rod-to-core) are connected by linker proteins.

Cyanobacteria, a monophyletic lineage of gram-negative oxygenic photosynthetic bacteria, thrive ubiquitously and inhabits various ecological zones, from aquatic environments such as lakes, rivers, and oceans to arid deserts, polar regions, and caves, and even in symbiosis with other organisms such as fungi, to form lichens ^6^. Physiologically, cyanobacteria can produce oxygen through photosynthesis; therefore, they can produce biomass using solar energy. Cyanobacteria have recently garnered attention for their potential as green cell factories for the CO_2_-neutral biosynthesis of various products ^7, 8^. To date, we have established CO_2_-derived useful material production systems, such as terpenoids and benzenoids, by metabolic engineering of the freshwater cyanobacterium *Synechococcus elongatus* PCC 7942 (hereafter *Synechococcus* 7942) ^9, 10^.

In their natural environment, cyanobacteria employ a distinct composition of disks within their PBS rods to optimize light harvesting based on available light wavelengths. For instance, freshwater cyanobacteria such as *Synechococcus* 7942 utilize only PCs bound to PCB to efficiently absorb the orange light ^11^. In contrast, PBS of cyanobacteria thriving in more deep-water environments has been optimized for light absorption in shorter wavelength light conditions, such as green light (GL), which can reach deep-water environments. The PBS rods in cyanobacteria such as *Synechococcus* sp. WH 7803 (*Synechococcus* 7803) contains PEs bound to phycoerythrobilin (PEB) along with PCs, allowing efficient light harvesting even in niche environments depleted of orange-to-red light ^12^.

PCB and PEB are isomers that differ only in the number of conjugated double bonds forming the chromophore and both are derived from a common biosynthetic precursor, namely biliverdin IXα, which is synthesized from heme (Figure 1). PCB:ferredoxin oxidoreductase (PcyA) synthesizes PCBs through a region-specific reduction in biliverdin IXα ^13^. In *Synechococcus* 7803, PEB is synthesized from biliverdin IXα via intermediate 15,16-dihydrobiliverdin (DHBV) by the sequential action of two reductases, namely DHBV:ferredoxin oxidoreductase (PebA) and PEB:ferredoxin oxidoreductase (PebB) (Figure 1) ^12, 13^. To form rod phycobiliproteins such as PCs and PEs, the synthesized PCBs and PEBs are covalently bound to specific apoproteins (phycocyanin alpha chain [CpcA] and phycocyanin beta chain [CpcB] for PC and phycoerythrin alpha chain [CpeA/MpeA] and phycoerythrin beta chain [CpeB/MpeB] for PE) using specialized lyases ^14-16^.

**Figure 1.**
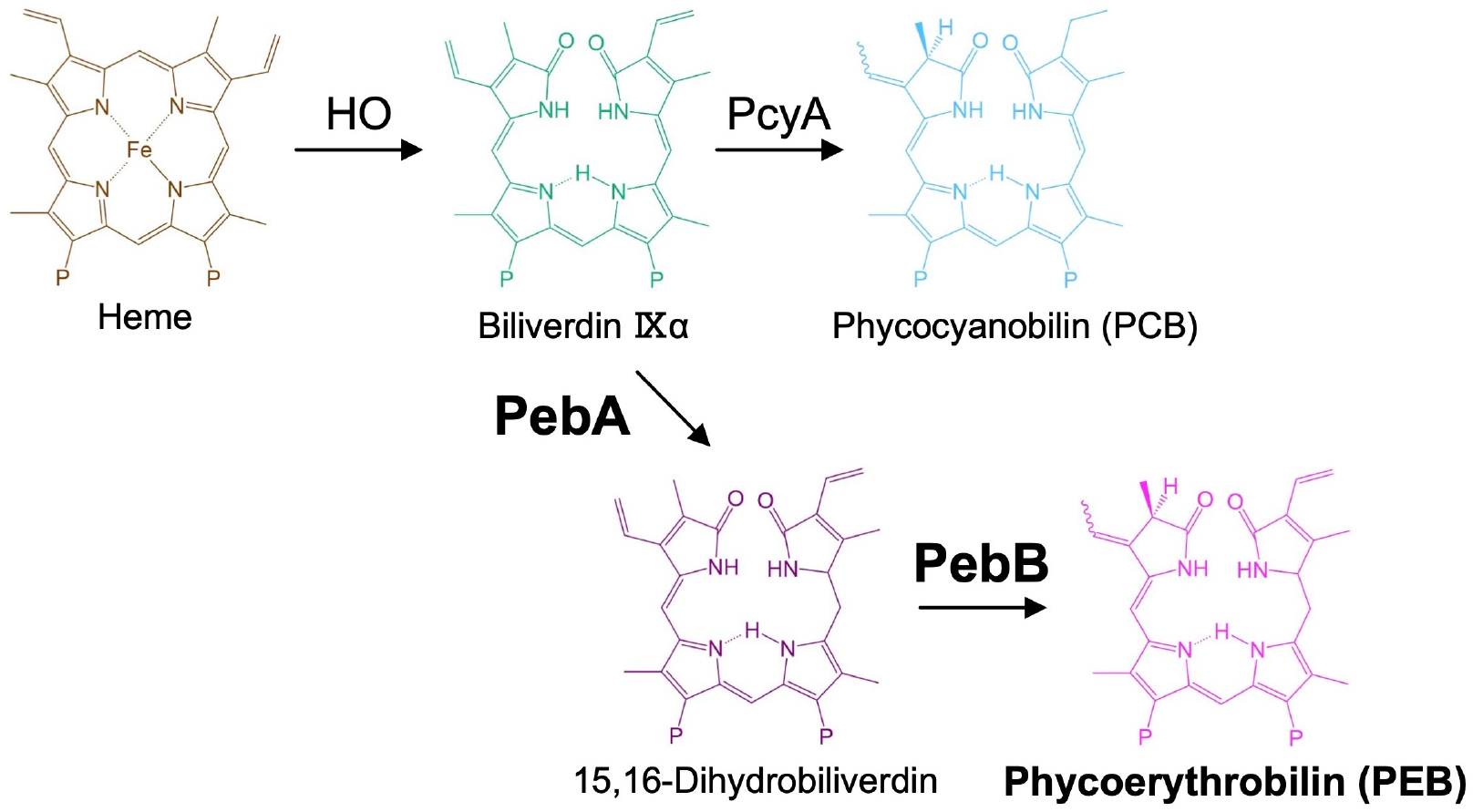
PCB and PEB metabolic pathways. Schematic overview of phycobilin biosynthesis from heme. In cyanobacterium, *Synechococcus* 7942, biliverdin IXα can be further converted to phycocyanobilin by the ferredoxin-dependent bilin reductase PcyA (top reaction). PCB, phycocyanobilin; PcyA, phycocyanobilin:ferredoxin oxidoreductase; PEB, Phycoerythrobilin; PebA, 15,16-dihydrobiliverdin:ferredoxin oxidoreductase; PebB, phycoerythrobilin:ferredoxin oxidoreductase; and HO, heme oxygenase.

Most PBS-related genes form functional clusters in the cyanobacterial genome, possibly because they have been acquired by horizontal gene transfer ^12^. Nevertheless, our understanding of the potential of PBS to accept heterogeneous components and their evolutionary capacity remains limited. A study showed that the constitutive co-expression of PebA and PebB (PebAB) in *Synechococcus* sp. PCC 7002 (*Synechococcus* 7002) resulted in cellular browning due to PEB accumulation ^17^. Interestingly, in *Synechococcus* 7002, PEB binds to endogenous CpcA without PEB lyase. However, it is genetically unstable, and the characteristics of PEB-bound PBS complexes remain largely unexplored.

Herein, we aimed to overcome these challenges and elucidate the functional implications and evolutionary importance of PEB-associated PBS complexes. We implemented a novel approach in the design of *Synechococcus* 7942 strains, enabling precise control of PEB metabolism. This engineered strain, *Synechococcus* 7942 PEB1, was examined for the effects of controlled PEB accumulation in the PBS complex. We examined if PEB1 exhibited any reversible color change owing to the precise control of PEB metabolism and depending on the induction level of PebAB. Additionally, the substitution of PCB with PEB and the disassembly of the PBS rod complex were investigated at high induction levels. Furthermore, the effect of low-level PebAB expression on the formation of full-size stable chimeric PBS complex and partial substitution of PCB with PEB. The results of this study may contribute to efficient light harvesting in the green spectrum and energy transfer to the photosynthetic reaction center.

## RESULTS AND DISCUSSION

### Construction of cyanobacterial strains with complete control over PEB levels

To fully regulate PEB metabolism, cyanobacterium *Synechococcus* 7942 was selected as the experimental organism; it can strictly regulate exogenous gene expression using isopropyl ß-D-1-thiogalactopyranoside (IPTG) ^9, 10^. *pebAB* of marine cyanobacterium *Synechococcus* 7803 was placed under the IPTG-inducible *spac* promoter and introduced into a neutral site (NS) on the chromosome of *Synechococcus* 7942, and the resulting strain was named *Synechococcus* 7942 PEB1 (Figure 2A). Brown colonies appeared on plates containing 100 μM IPTG under the white light (WL) irradiation, whereas green color was retained in the medium without IPTG and in the control strain without *pebAB* (Figure 2B). In contrast to the previous study on *Synechococcus* 7002 ^17^, no green suppressor colonies appeared in the PEB1 strain after >10 days of incubation with IPTG (Figure 2B), indicating that the PEB1 strain can stably control the PEB metabolism. In the liquid medium, browning was observed in an IPTG-dose-dependent manner (Figure 2C). The absorption spectra of these cultures showed a decrease in the 625-nm peak representing PC and an increase in the 550-nm peak following IPTG addition (Figure 2D). Because PEB can bind to the PC subunit CpcA and its complex exhibits an absorption peak similar to that of PE ^17^, the 550-nm peak observed in PEB1 was also considered to be similar to that of PEB-bound chimeric phycobiliprotein.

**Figure 2.**
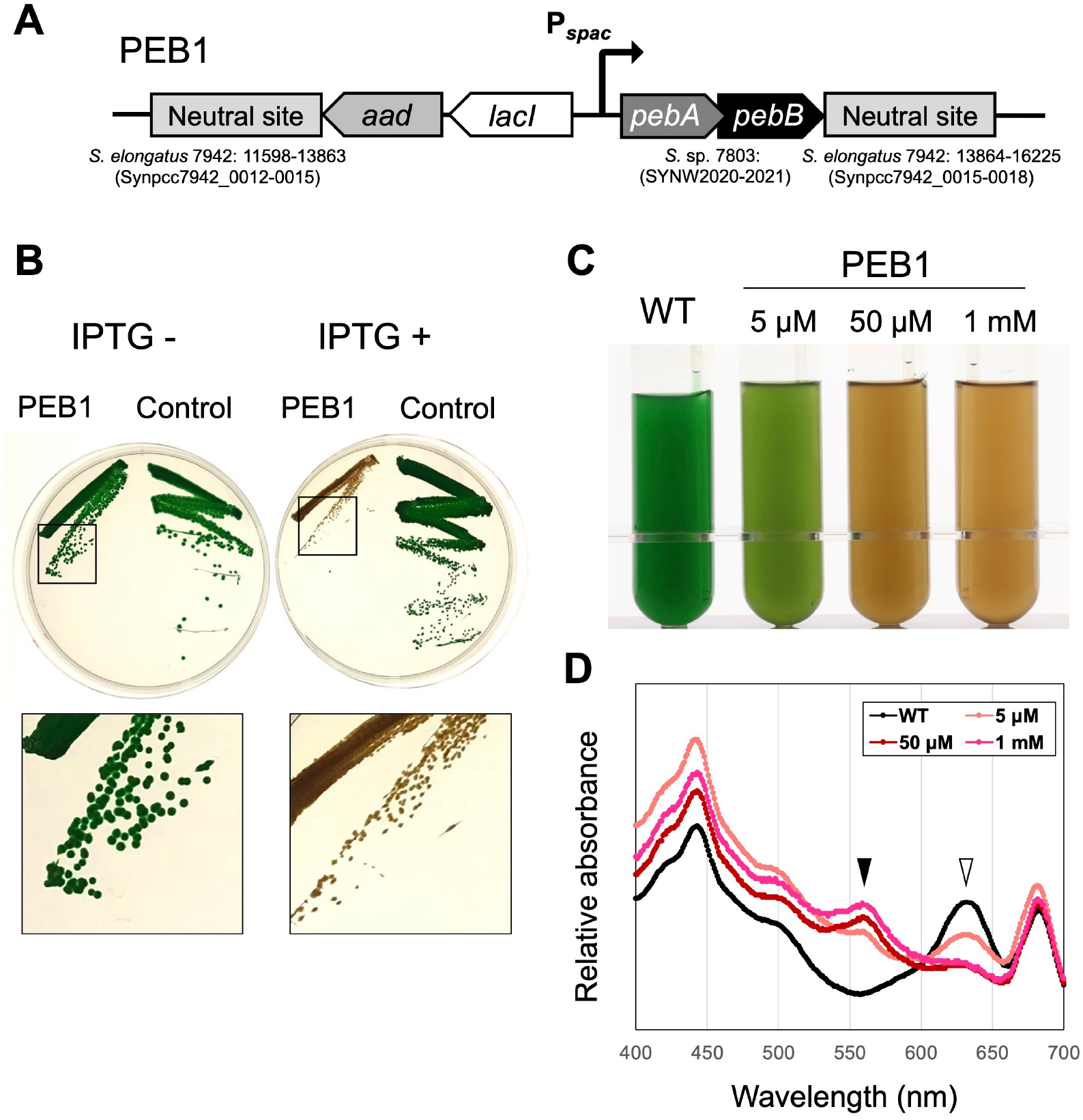
Construction of a PEB1 strain allowing control of PEB levels. (A) Schematic diagram of the genomic construct of *Synechococcus* 7942 PEB1 strain expressing *pebA* and *pebB* in an IPTG-dependent manner. The *pebA* and *pebB* genes of *Synechococcus* 7803 were placed under the IPTG-dependent *spac* promoter and introduced into the neutral site of *Synechococcus* 7942 along with the spectinomycin resistance gene (*aad*) and *lacI*. (B) The phenotype of the PEB1 strain on the plates. PEB1 and control strain transformed using the empty vector were grown for 10 days under the white light (60 μmol photons m^−2^·s^−1^) on the BG-11 plates containing 40 μg/mL spectinomycin with (right) and without 100 μM IPTG (left). Enlarged colony images are shown below. (C) Wild-type and PEB1 strains grown in the liquid medium under the white light. The strains were cultured in liquid BG11 in a photobioreactor at 30°C, with air bubbling for 1 week. (D) Absorption spectrum of the cultures. The spectra were normalized at 680 nm. Closed arrowhead: peak at 560 nm and open arrowhead: peak at 625 nm. PEB, phycoerythrobilin; IPTG, isopropyl ß-D-1-thiogalactopyranoside; and WT, wild type.

Our further studies revealed that PEB1 could dramatically and reversibly control PEB levels. Brown PEB1 cells were cultured in a liquid medium containing 1 mM IPTG for 1 week and then used for subsequent experiments. When brown cells were further cultured with IPTG under WL irradiation, the cell color changed to pink, whereas the color recovered from brown to green in the medium without IPTG (Figure 3A; #3 and #4) (Supplementary Movie). Along with the color change, the absorbance spectra and growth of PEB1 cells changed with the addition of IPTG (Figures 3B and 3C). In pink cells, the absorption peak at 550 nm was notably higher than those in brown and green cells, and growth inhibition was observed under WL (Figure 3B). These results suggest that pink cells accumulated large amounts of PEB; however, its utilization for light harvesting under WL conditions was limited.

**Figure 3.**
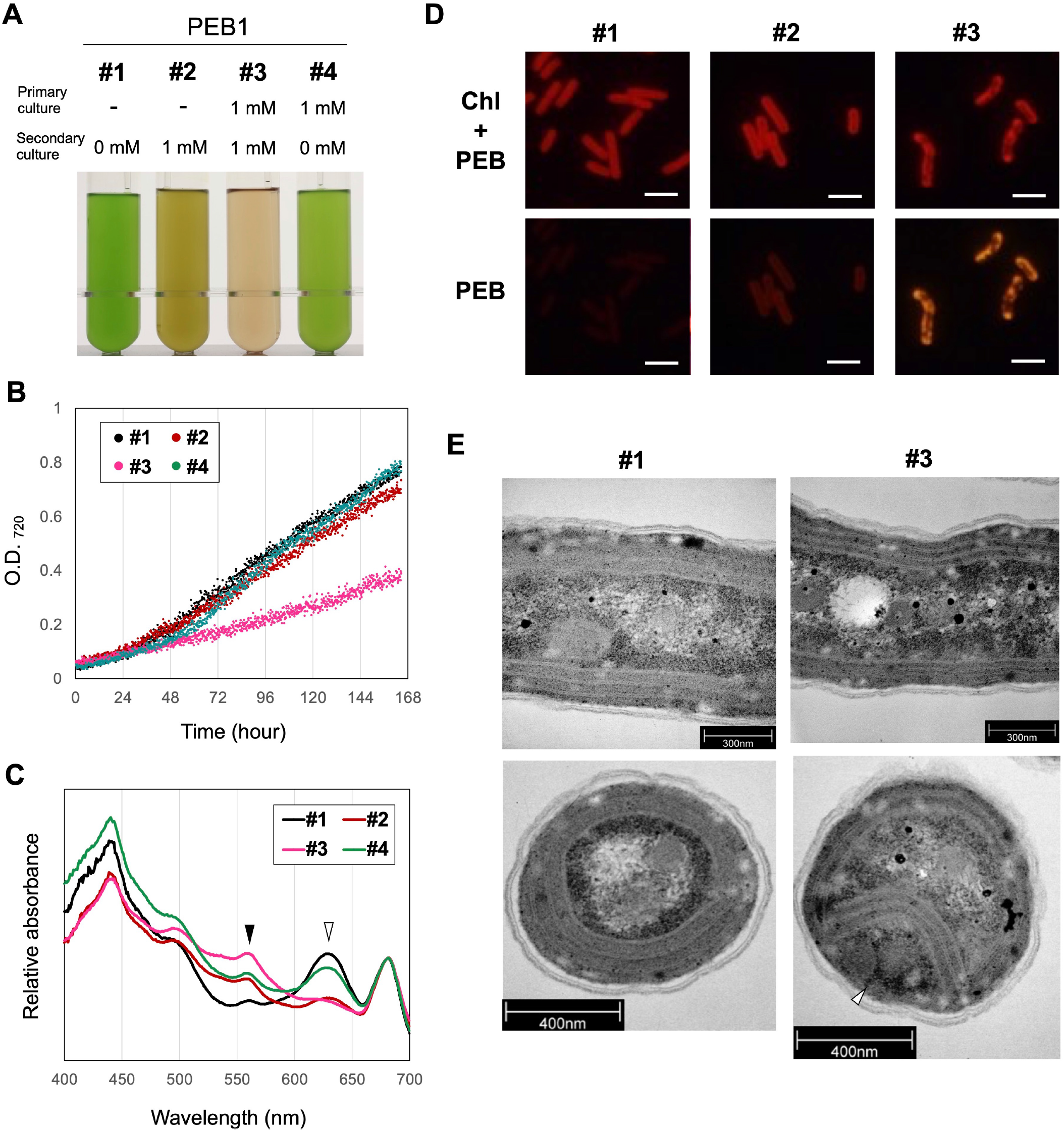
Effects of PEB hyperaccumulation and reversible phenotypic change in the PEB1 strain. The PEB1 culture that turned brown after growing in the BG11 medium containing 1 mM isopropyl ß-D-1-thiogalactopyranoside (IPTG) for 1 week under white light (#2) was additionally cultivated with (#3) and without IPTG (#4) for another week and was compared to the culture grown without IPTG (#1). (A) PEB1 cultures after 1-week incubation in the liquid medium. (B) Growth curves of PEB1 cultures. (C) Absorption spectra of the cultures. Closed arrowhead: peak at 560 nm and white arrowhead: peak at 625 nm. The spectra were normalized at 680 nm. (D) Fluorescence microscopy of Chl and PEB fluorescence in #1–3 cells. White bar: 5 μm. (E) Transmission electron microscopy of #1 and #3 PEB1 cells. Upper panels, horizontal images, lower panels, and vertical images. The white arrowhead indicates the space and unusual structure observed between the cell and the thinned thylakoid membrane. PEB, phycoerythrobilin; Chl, chlorophyll; and O.D., optical density.

Microscopic observations revealed abnormal cell morphology owing to PEB accumulation. In brown cells, PEB and chlorophyll fluorescence were distributed throughout the cells, whereas in pink cells, foci of strong PEB fluorescence were observed with an abnormal distribution of chlorophyll (Figure 3D). Transmission electron microscopy (TEM) results revealed that the pink cells had an abnormal cell morphology, with narrow intervals between thylakoid membranes (Figure 3E), suggesting that the amount of PBS complexes was reduced at these intervals. Additionally, unusual structures were observed in the space between the cell and the thylakoid membrane in pink cells (Figure 3E, white arrowhead). These unique structures possibly arise from an overproduction of PEB-bound chimeric phycobiliproteins.

### Collapse of PBS rod complexes because of excessive PEB accumulation

We examined the sizes and properties of PBS complexes in the PEB1 strain. To fractionate the full-size PBS complex and separate its components, cell extracts of PEB1 were subjected to sucrose density gradient (SDG) centrifugation. In the brown and pink cells, the fraction containing full-size PBS complexes observed in the bottom layer of the gradient disappeared. A red-colored fraction was observed in the upper layer (Figure 4A; #2 and #3). Absorption spectra with a peak at 550 nm were observed in these red-colored upper fractions (Figure 4B), indicating that, in the brown and pink cells (Figure 2A; #2 and #3), the PBS rod complexes were disintegrated due to excessive PEB accumulation.

**Figure 4.**
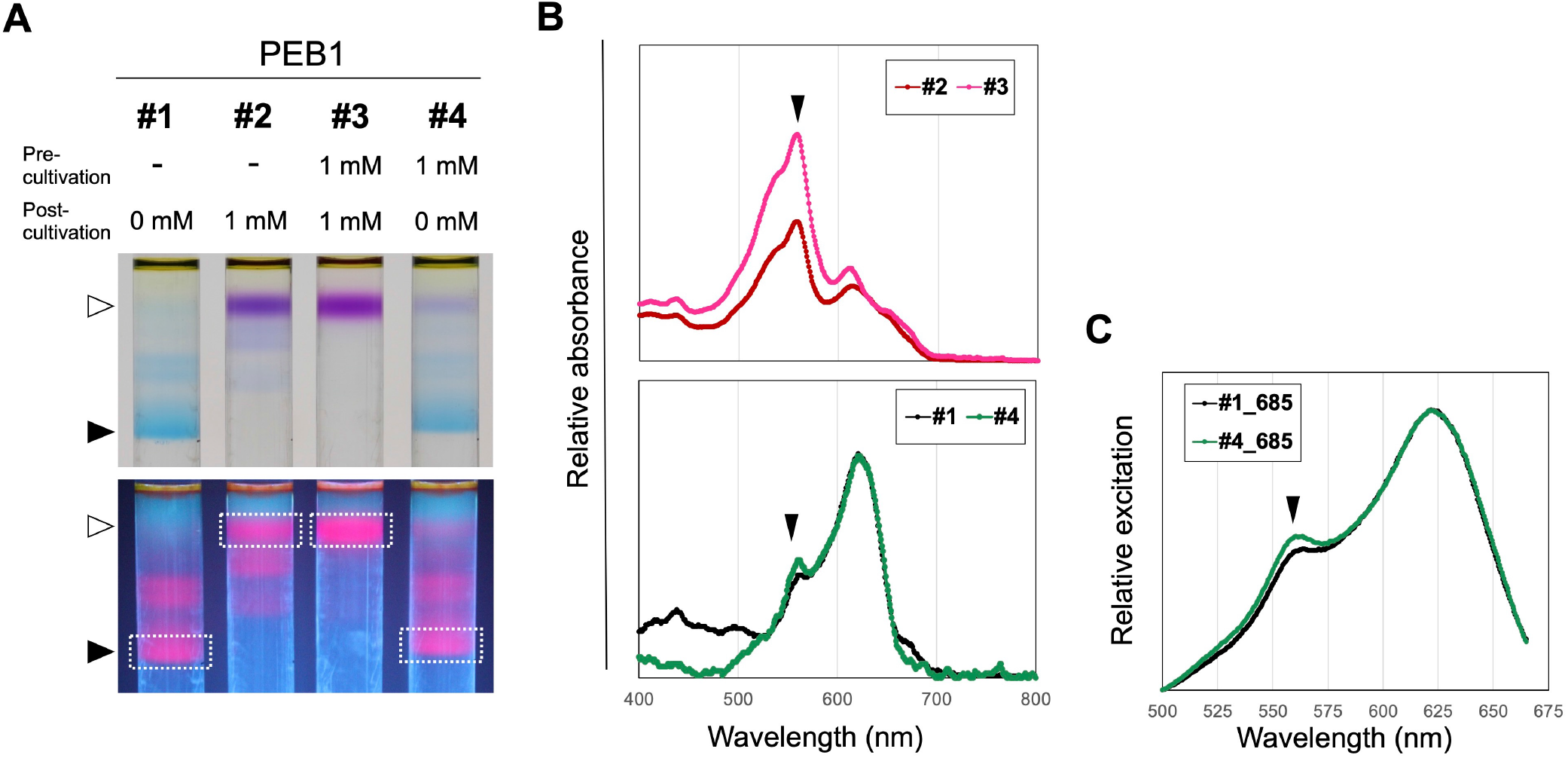
Comparison of the properties of phycobilisome (PBS) complexes fractionated by sucrose density gradient (SDG) centrifugation. PBS was fractionated from the cell extracts of PEB1 cultures (A) PBS fraction images obtained after SDG centrifugation. The lower image shows the excitation of phycobiliproteins after ultraviolet-A irradiation. The fractions subjected to absorption spectroscopy (B) and excitation spectroscopy (C) are enclosed by dotted lines. (B) Absorption spectra of phycobiliprotein fractions. The samples indicated by open and closed arrowheads in (A) are shown in the upper and lower panels, respectively. The spectra were normalized at 630 nm. (C) Excitation spectra of the PBS fractions emitting allophycocyanin fluorescence (685 nm). Spectra of the bottom fraction containing mature PBS (closed arrowheads in A) normalized at 630 nm. Closed arrowheads: peak at 560 nm. PEB, phycoerythrobilin.

When incubated without IPTG, the size and color of the PBS complexes in brown cells recovered to their original state (Figures 4A and 4B; #1 and #4). A small peak at 550 nm was detected in the absorbance spectra of full-size PBS complexes in these cultures (Figure 4B; #1 and #4). The excitation spectra at 685-nm fluorescence emitted by APC in PBS cores showed that the green PEB1 strains (Figure 2A; #1 and #4) had a peak near the absorption wavelength of PE and PC (560 nm and 620 nm, respectively) (Figure 4C). These results indicated that the PEB1 strain accumulated a small amount of PEB in the PBS full complex, even without IPTG, and that PBS can absorb short-wavelength GL through PEB and transfer energy to the PBS core.

Reportedly, PEB can bind to CpcA^17, 18^; similar bindings between PEB and PBS proteins were expected in the PEB1 strain. The color of the phycobiliprotein extracts obtained from PEB1 was different from that of the wild-type (WT) (Figure 5A), and red bands were observed in both the lower and higher bands when separated by sodium dodecyl sulfate (SDS)-polyacrylamide gel electrophoresis (PAGE) (Figure 5B). Proteins were extracted from each band and the absorption spectra were compared. The absorption wavelength of the lower band shifted to a shorter wavelength in the PEB1 strain (Figure 5C), suggesting preferentialinteraction between apoproteins of this size (such as CpcA) and PEB rather than PCBs. *Synechococcus* 7002, *Synechocystis* sp. PCC 6803 (*Synechocystis* 6803) and *Synechococcus* 7942, which were suggested to interact with PEB and CpcA^17, 18^, all utilize the E/F-type bilin lyases (CpcE and CpcF), for the binding of PCBs to apoproteins ^19^. These observations indicate that PEB is incorporated into CpcA by an E/F-type lyase, consistent with previous study *in vitro*^19^. In the higher fraction containing CpcB, additional peaks were detected in the absorption spectrum of PEB1. In both IPTG conditions, peaks at shorter wavelengths were observed (Figure 5C), whereas a further peak appeared on the longer-wavelength side under 1 mM IPTG supplementation conditions (Figure 5C, arrowhead). These results suggest that this fraction contained several types of phycobiliproteins, and that CpcB must also be able to bind to PEB. Although binding of PEB to CpcB and the PBS core may have caused the PBS disassembly, further studies are required to elucidate the reasons for this. A similar observation was reported for the binding of PEB to APC in *Synechocystis* 6803 strain lacking PC ^20^; APC naturally forms trimeric aggregates when PCBs bind, but substitution with PEB inhibited the next stage of monomer aggregation.

**Figure 5.**
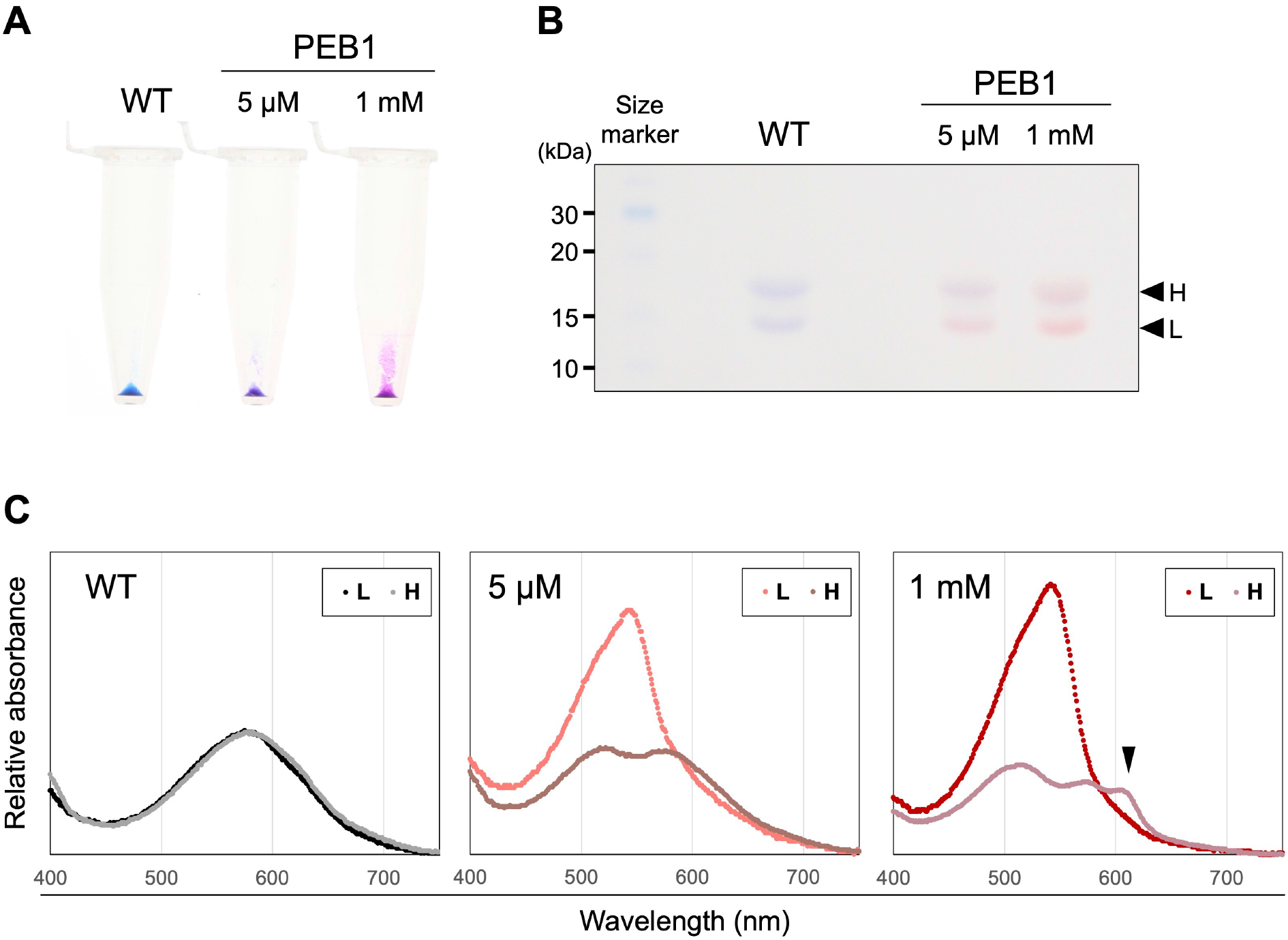
PEB preferentially binds to the C-phycocyanin alpha chain (CpcA) WT and PEB1 strains were grown under white light conditions in a liquid BG-11 medium. The indicated amounts of isopropyl ß-D-1-thiogalactopyranoside were added to the culture. (A) Pellet images of phycobiliproteins extracted from the WT and PEB1 cells. (B) Coomassie brilliant blue-stained gel images of phycobiliproteins. Arrowheads indicate phycobiliproteins with lower (L) and higher (H) molecular weight, containing phycocyanin and allophycocyanin subunit proteins, respectively. (C) Absorption spectra of phycobiliproteins -extracted gel. Each set of spectra was normalized to 584 nm. PEB, phycoerythrobilin, and WT, wild-type.

### Establishment of GL-utilizing PBS by controlling PEB levels

Because the PEB1 strain was expected to form full-size PEB-bound PBS complexes at low levels of *pebAB* induction, we further investigated the utilization of GL by these PBS complexes. Full-size PBS complexes were isolated from the PEB1 strain with low induction of *pebAB* by adding 5 μM IPTG. The fluorescence excitation spectrum of the PBS complex revealed an additional peak at 560 nm, which was not observed for the WT strain (Figures 6A and 6B). Growth tests using a photobioreactor under GL (530 nm) showed that the PEB1 strain (supplemented with 5 μM IPTG) grew considerably faster than the WT strain with increasing light intensity (Figures 6C and 6D). The energy transfer of the PEB1 culture using the 77K fluorescence spectra indicated that PEB1 could utilize GL. The fluorescence emitted by photosystem (PS)II (695 nm) and PSI (715 nm) was observed following excitation at 530 nm (Figure 6E). Furthermore, the 77K excitation spectrum of PSII (695 nm) showed peaks corresponding to the absorption of PEB, PC, and APC at 560 nm, 630 nm, and 650 nm, respectively, indicating the ability of PEB1 to transfer GL energy to PSII (Figure 6F).

**Figure 6.**
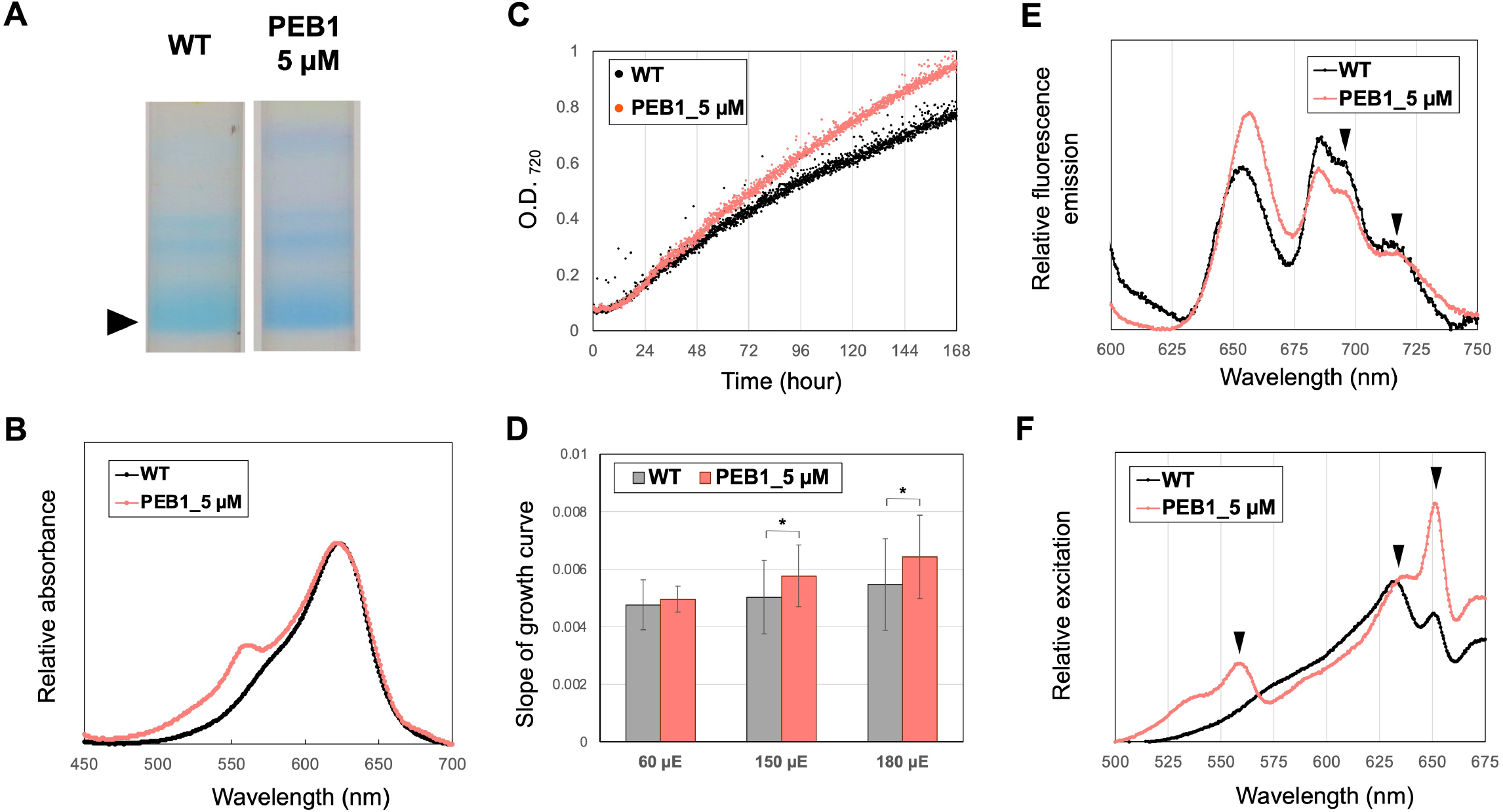
Green light (GL) adaptation of PEB1. (A) Phycobilisome fraction images after sucrose density gradient centrifugation. The samples were prepared from the cells grown under the white light. Isopropyl ß-D-1-thiogalactopyranoside (5 μM) was supplied to PEB1 cultures to express the appropriate amount of 15,16-dihydrobiliverdin:ferredoxin oxidoreductase (PebA) and PEB:ferredoxin oxidoreductase (PebB). (B) Absorption spectra of the bottom fractions indicated by closed arrowhead in (A). (C and D) Comparison of growth under the GL irradiation (180 μmol photons m^−2^·s^−1^). (C) Growth curves of WT and PEB1 cultures. (D) Comparison of growth rate estimated based on the slopes of the growth curves. The intensity of irradiated GL is shown on the x-axis. For statistical evaluation, *p*-values were calculated using the paired *t*-test in Microsoft Excel, \**p* < 0.05. (E and F) The 77K fluorescence and excitation spectra of WT and PEB1 cells. (E) Fluorescence spectra were excited at a wavelength of 530 nm. The spectra were normalized at 720 nm. The peaks of fluorescence emitted by photosystem (PS) II (695 nm) and PSI (715 nm) are indicated by arrow heads. (F) Excitation spectra measuring fluorescence at 695 nm. The spectra were normalized at 625 nm. The peaks of the absorption of 560 nm, 630 nm, and 650 nm are indicated by arrow heads. PEB, phycoerythrobilin and WT, wild-type.

### Growth advantage of PEB1 strain under GL

To examine the growth advantages of PEB1 strain under GL, we performed a competition assay. For the competitor strain for PEB1, we constructed a *Synechococcus* 7942 CAT strain carrying the chloramphenicol acetyltransferase gene (*cat*) at an NS on the chromosome (Figure 7A). The CAT strain was co-inoculated with the PEB1 strain to the BG-11 liquid medium containing 5 μM IPTG with no antibiotics and cultured under the GL condition. After 1 and 2 weeks, the culture was spread onto BG-11 medium plates containing spectinomycin or chloramphenicol, and the population of each strain was calculated based on the number of colonies on the plates. The results showed a significant increase in the population of the PEB1 strain compared with that of the CAT strain at both 1 and 2 weeks (Figure 7B), indicating that the PEB1 strain preferentially grew under GL compared with the CAT strain.

**Figure 7.**
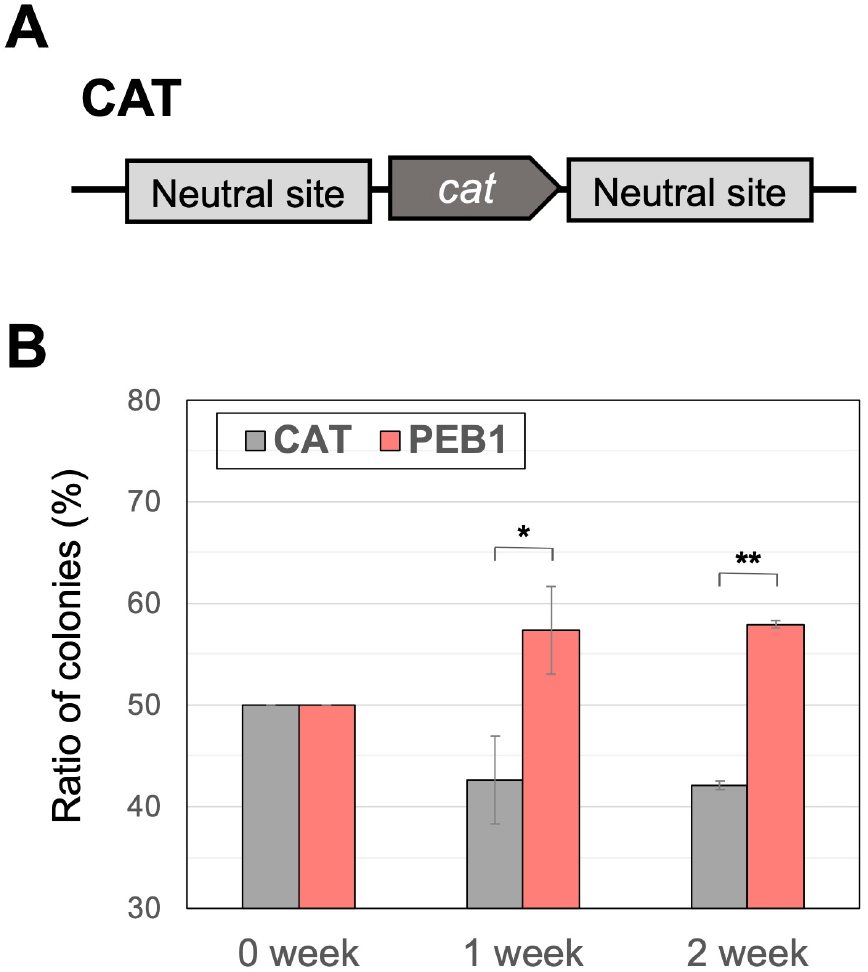
Competition assay. (A) Schematic diagram of the genomic construct of CAT strain. (B) The populations of CAT and PEB1 strains. PEB1 was grown together with CAT under green light irradiation (180 μmol photons m^−2^·s^−1^). Before and after the incubation, the culture was spread on the BG-11 plates containing spectinomycin or chloramphenicol and incubated for 1 week. The populations of CAT and PEB1 strains were estimated based on the number of colonies appearing on the plates. For statistical evaluation, *p*-values were calculated using the paired *t*-test in Microsoft Excel, \**p* < 0.05; \*\**p* < 0.01. CAT, *Synechococcus* 7942 CAT strain and PEB, phycoerythrobilin.

## CONCLUSIONS

To conclude, we established a precise and stable control of PEB levels in *Synechococcus* 7942 by controlling the expression of *pebAB* and showed the effect of PEB in PCB-bound PBS. The proposed model for the effects of PEB production is illustrated in Figure 8. The *Synechococcus* 7942 WT strain, which uses PCBs in PBS, efficiently transferred red light to the PS. In the PEB1 strain, at high induction levels of PEB, PBS disassembly occurred, whereas at low levels of induction, PBS was constructed to absorb not only red light but also GL, and it could transfer the light energy to the PSs.

**Figure 8.**
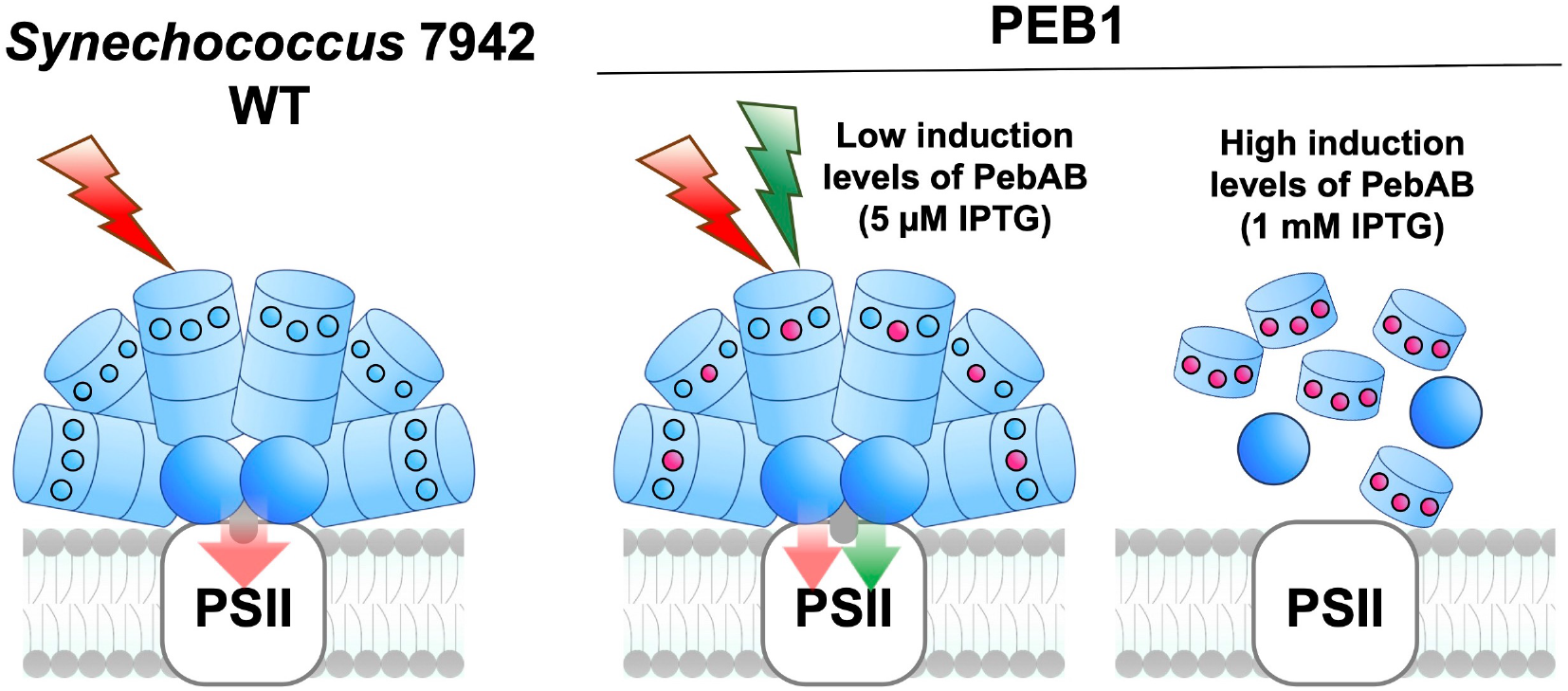
Hypothetical model of phycobilisome (PBS) during the induction of PEB. Hypothetical model of PBS in *Synechococcus* 7942 WT and PEB1 at high and low PEB induction levels Only bilins (namely PCB and PEB) that bind to the outer disk are shown in blue (PCB) or red circles (PEB). PEB, phycoerythrobilin; WT, wild-type; PebAB, 15,16-dihydrobiliverdin:ferredoxin oxidoreductase and PEB:ferredoxin oxidoreductase; IPTG, isopropyl ß-D-1-thiogalactopyranoside; and PS, photosystem.

In nature, several cyanobacteria have been found to flexibly and dynamically rearrange their photosynthetic apparatus, including PBS, in response to light environment ^21^. These changes are accomplished through a complex cellular response involving many genes to adapt PBS to the light environment, including signaling systems, bilin synthases, apoproteins, and lyases that catalyze the binding of pigments to proteins. Herein, we have shown that it is possible to artificially alter the properties of PBS by manipulating the PEB metabolism. This is the first study to construct a cyanobacteria that can utilize GL for faster growth by genetic modification. The accumulation of a small amount of PEB-based PCs under precise control is the key to this achievement. The expression of additional factors such as PEB-binding apoproteins and other components of PEB-PBS may allow for the breeding of cyanobacteria with PBS, which is more suitable for various light conditions.

Our study provides an experimental basis for future studies to elucidate the underlying mechanisms governing PBS adaptation, and it offers potential applications in bioengineering, synthetic biology, and renewable energy research. Recent attempts to modify bilin metabolism in *Synechocystis* 6803 highlight the need to understand the unique PBS regulation across diverse cyanobacteria ^18^. This research can deepen our understanding and aid in the development of artificial pigments. Additionally, our research stands at the forefront of advancing the development of artificial pigments. Notably, our study on PEB-based PCs, which are powerful fluorescent emitters, holds the potential to revolutionize industrial applications using novel fluorescent reagents. In particular, the higher accumulation of small pink-colored components under higher IPTG concentrations (Figure 4A; 2# and 3#) would be advantageous for the production of the fluorescent reagents. Unlike natural PE synthetic cyanobacteria, model cyanobacteria, such as *Synechococcus* 7942, offer faster growth rates and easier manipulation. Hence, this pioneering synthetic pathway holds immense promise for significantly enhancing phycobiliprotein production.

## METHODS

### Culture conditions for cyanobacteria

Cyanobacterium *Synechococcus* 7942 WT strain and its derivatives were grown photoautotrophically at 30°C under continuous WL illumination (60 μmol photon m^−2^·s^−1^) in the modified BG-11 medium, which contained double the usual amount of sodium nitrate (final concentration = 35.3 mM) and 20 mM 4-(2-hydroxyethyl)-1-piperazineethanesulfonic acid (HEPES)–KOH (pH 8.2) with 2% CO_2_ bubbling. When appropriate, spectinomycin (final concentration = 40 μg/mL) and chloramphenicol (final concentration = 10 μg/mL) were added to the media.

The growth experiments were performed using a Multi-Cultivator MC1000-OD (Photon Systems Instruments, Czech Republic) under continuous illumination with WL (60 μmol photons m^−2^·s^−1^) or GL (530 nm; 60, 150, and 180 μmol photons m^−2^·s^−1^), with bubbling ambient air. After preincubation on the BG-11 medium plate for 1 week, cells were harvested, inoculated into liquid BG-11 medium with IPTG at an optical density (OD)_720_ of 0.05, and cultivated for 1 week. An approximate growth curve was constructed based on the OD values up to day 7 (168 h). Measurements were conducted in triplicate, and the slope of the approximate growth curve was compared with the growth rate. Significance was determined by paired *t*-test in Microsoft Excel.

### Strain construction

The DNA fragments of PEB synthase genes *pebA* and *pebB* from *Synechococcus* 7803 (open reading frame ID: SYNW2020 and 2021), upstream and downstream of the NS of *Synechococcus* 7942 genomic region, and spectinomycin-resistant gene with *lacI–spac* promoter unit were amplified through polymerase chain reaction (PCR) using PrimeSTAR DNA polymerase (TaKaRa, Shiga, Japan), along with appropriate primer sets (namely F1/R2, F3/R4, F5/R6, F7/R8, and F9/R10; Table S1). In total, five DNA fragments were recombined through PCR using the primer set F3/R6 (Table S1), and the resulting fragment was introduced into *Synechococcus* 7942 to obtain spectinomycin-resistant transformants. Successful transformation of the strains with the DNA fragments was confirmed by PCR amplification using the specific primer set F11/R12, and the resulting strain was named *Synechococcus* 7942 PEB1.

The competitor strain *Synechococcus* 7942 CAT, containing *cat* at the NS, was constructed and used for the growth competition test under GL. In total, three DNA fragments containing *cat*, upstream and downstream of NS, were amplified using appropriate primer sets (namely F13/R14, F3/R15, and F16/R6; Table S1) and recombined by PCR using the primer set F3/R6. The resulting fragment was introduced into *Synechococcus* 7942 to obtain chloramphenicol-resistant transformants. Successful transformation of the strains was confirmed by PCR using the primer set F11/R12.

### Fluorescence microscopy

Fluorescent images were obtained using a BX53 microscope (OLYMPUS, Tokyo, Japan) at 100× magnification with a DP71 digital camera (OLYMPUS) and the DP Controller software ver. 3. 3. 1. 292 (OLYMPUS). Chlorophyll and PE autofluorescence were observed using the U-MWIG3 and U-FRFP filter units (OLYMPUS), respectively.

### TEM

PEB1 strain was inoculated in the BG11 liquid medium at an OD of 0.1 with and without 1 mM IPTG and incubated for 24 hours at 30°C. The cells were collected by centrifugation at 3,000× *g* for 10 minutes at 25°C and fixed with 1 mL of ice-cold 2% glutaraldehyde solution overnight at 4°C (prefixation). The cells were collected by centrifugation and washed with 1 mL of 0.1 M phosphate buffer (pH 7.4) overnight at 4°C. For postfixation, ice-cold 2% osmium tetroxide was added to the samples and incubated for 3 hours at 4°C. The stained cells were dehydrated with increasing concentrations of ethanol (50%, 70%, 90%, and 100%) for 15 min each, and the dehydrated cells were embedded in a gelatin capsule with epoxy resin for 2 days at 60°C. Ultra-thin sections (80–90 nm), which were stained with 2% uranyl acetate for 15 minutes and lead staining solution for 2 minutes, were subjected to TEM (JEM1200EX; JEOL, Tokyo, Japan) at the Hanaichi Ultra Structure Research Institute (Aichi, Japan).

### Analysis of phycobiliproteins

The cells were harvested and resuspended in 100 μL of A buffer (containing 10% glycerol, 100 mM NaCl, 20 mM HEPES–NaOH; pH 7.5) and then disrupted with glass beads using a bead beater (Micro Smash MS-100R, TOMY Seiko Co., Tokyo, Japan). After the centrifugation at 20,000× *g* for 1 minute at 18°C, 60 μL of supernatants were moved to fresh tubes and mixed with 240 μL of acetone (final concentration = 80%) to remove chlorophyll and carotenoids from the cell lysates. The resulting samples were centrifuged at 20,000× *g* for 1 minute and pellets containing phycobiliproteins were obtained. Bilins bindings to apoproteins were analyzed by SDS-PAGE. Next, 20 μL of supernatants mixed with loading dye (final concentration = 0.0625 Tris-HCl, 10% glycerol, 2% SDS, and 0.01% bromophenol blue) were separated on a 15% (w/v) polyacrylamide gel. After PAGE, the excised gels were extracted using ATTOPREP MF (ATTO, Tokyo, Japan), and the absorption spectra were measured.

### Isolation of PBSs by SDG centrifugation

The *Synechococcus* 7942 culture (OD = 1.0; 30 mL) was harvested by centrifugation (3,000× *g*, 10 minutes, 25°C). After washing with 1 mL of 0.6 M potassium phosphate (KP) buffer (pH = 7.0), the cells were again centrifuged (3,000× *g*, 10 minutes, 25°C) and were stocked in a freezer until used for analysis. The cells were washed twice in 0.6 M KP buffer and resuspended in 0.6 mL of 0.6 M KP. Cells were lysed by vortexing with glass beads and PBS complexes were extracted from the thylakoid membranes by vortexing with Triton X-100 (final concentration = 2%) for 15 minutes. After centrifugation (20,000× *g*, 20 minutes, 18°C), 200 μL of supernatant was loaded onto sucrose gradients (10–50% sucrose in 0.6 M KP buffer) prepared in tubes (14 × 89 mm, Open-Top Thinwall Ultraclear tubes [Beckman Coulter, CA, USA]) using a Gradient Master (Beckman Coulter). The gradients were centrifuged at 154,300× *g* for 18 hours at 18°C (SW41Ti rotor, Optima XE-90 Ultracentrifuge [Beckman Coulter]). After centrifugation, the samples were irradiated with ultraviolet (UV)-A (365 nm) using a UV lamp (UVP Inc.) to observe the PC and PE fluorescence of phycobiliproteins.

### Spectrometry

The absorption spectra of the cells, phycobiliproteins, and PBS complexes were measured at 25°C using a spectrophotometer (model UV-1800, Shimadzu, Japan). Excitation spectra were recorded at 685 nm (APC) using a spectrophotometer (model FP-8200, JASCO, Japan). To measure the 77K fluorescence excitation spectra, the samples were frozen in liquid nitrogen and PSII fluorescence (695 nm) was measured (model RF-6000, Shimadzu, Japan).

### Growth competition test under GL

After preincubation on BG-11 medium plates containing spectinomycin (for PEB1) or chloramphenicol (for CAT), both PEB1 and CAT cultures were inoculated in the BG11 liquid medium containing 5 μM IPTG with no antibiotics at an OD of 0.025 each and incubated for 2 weeks at 30°C under continuous GL (530 nm, 180 μmol photons m^−2^·s^−1^) with bubbling ambient air. Immediately after inoculation, 1 week, and 2 weeks later, 10 mL of culture medium was incubated on plates containing spectinomycin or chloramphenicol, and the number of colonies that grew on each plate after 1-week incubation was counted. Measurements were conducted in triplicate, and the data are presented as mean ± standard deviation (SD). Significance was determined by paired *t*-test in Microsoft Excel.

## ASSOCIATED CONTENT

Color change of PEB1 cultures over time (Movie) and lists of primers (Table S1)

## AUTHOR INFORMATION

### Author contributions

R.N. and S.W. designed the concept and the experiments of this study; M.S., T.K., K.M. and M.W. performed the experiments; M.W., R.N., M.I. and S.W. analyzed the data; M.W., R.N., M.I. and S.W. wrote the manuscript.

## ACKNOWLEDGMENTS

We are grateful to Prof. Mitsuhiro Itaya and Prof. Taku Chibazakura for their valuable comments on the concept of this study. We thank Prof. Fujio Kawamura and Dr. Claudia Steglich for providing DNA sources of antibiotic resistant genes and *Synechococcus* 7803 genome, respectively. We thank Dr. Keita Miyake and Dr. Hiroki Hoshino for their assistance with spectrometry. This work was supported by the Ministry of Education, Culture, Sports, Science and Technology of Japan to SW (20K05793 and 23H02130) and the Advanced Low Carbon Technology Research and Development Program (ALCA) of the Japan Science and Technology Agency (JST), New Energy and Industrial Technology Development Organization (NEDO, JPNP17005) (to S.W.).

## ABBREVIATIONS

PBS: phycobilisome
PC: phycocyanin
PE: phycoerythrin
PEC: phycoerythrocyanin
APC: allophycocyanin
PCB: phycocyanobilin
PEB: phycoerythrobilin
SDG: sucrose density gradient

