## Supplementary material for "Functional Modification of Cyanobacterial Phycobiliprotein and Phycobilisomes through Bilin Metabolism Control": Supplementary Materials_240103.pdf

### 1 Movie：Color change of PEB1 cultures over time (mp4. file)

To prepare the primary cultures, *Synechococcus* 7942 PEB1 strain harvested from BG11 plates were inoculated into BG11 liquid medium with and without 1 mM IPTG and incubated for 3 days. Those cultures were further transferred to medium with and without 1 mM IPTG at OD=0.2 as secondary cultures and monitored for color change over 36 hours. The movie, compressed to one second per hour, shows 36 hours of recording.

### 2 Table

**Table S1. Oligonucleotide primers used in this study.**

| Primer name | Sequence (5′ to 3′)[a] |
|---|---|
| *Strain construction* | |
| F1 | CTTCAATCACTAGAGCTACGACAAGGAAGTTGAGGCGGC |
| R2 | GAGGAGAAATTAACTATGTTCGACTCTTTCCTCAAC |
| F3 | CGCGAATCCAGAACTTTGATGTCC |
| R4 | ACTTCCTTGTCGTAGCTCTAGTGATTGAAGGGGCCTGC |
| F5 | CTTCTCTCAATTAGCTCCACCGATGTAGCGGTC |
| R6 | CGATCGTCAAAGGTGAGTAGCCG |
| F7 | GAAAGAGTCGAACATAGTTAATTTCTCCTCTTTAATGAATTCAA |
| R8 | CTGCGCTTTTTTTCATCACTGCCCGCTTTCCAGTCGG |
| F9 | GAAAGCGGGCAGTGATGAAAAAAAGCGCAGCTGAAATAG |
| R10 | GCTACATCGGTGGAGCTAATTGAGAGAAGTTTCTATAGAATTTTTC |
| F11 | GAAGTGGCGATCGCCGTGATCG |
| R12 | GATCAACACGGTGCAGGGTGG |
| F13 | CTTCAATCACTAGAGCTGCGTTAGTCGTCATTAAGCA |
| R14 | GCTACATCGGTGGAGTTATAAAAGCCAGTCATTAGGCCTATCTGAC |
| R15 | TGACGACTAACGCAGCTCTAGTGATTGAAGGGGCCTGC |
| F16 | GACTGGCTTTTATAACTCCACCGATGTAGCGGTC |
